# A general strategy for generating expert-guided, simplified views of ontologies

**DOI:** 10.1101/2024.12.13.628309

**Authors:** Anita R. Caron, Aleix Puig-Barbe, Ellen M. Quardokus, James P. Balhoff, Jasmine Belfiore, Nana-Jane Chipampe, Josef Hardi, Bruce W. Herr, Huseyin Kir, Paola Roncaglia, Mark A. Musen, Helen Parkinson, James A. McLaughlin, Katy Börner, David Osumi-Sutherland

## Abstract

Annotation of biomedical entities with widely used, well-structured ontologies and ontology-aware tools ensures data and analyses are Findable, Accessible, Interoperable, and Reusable (FAIR). Standardized terms with synonyms support lexical search, while ontology structure enables biologically meaningful grouping of annotations, such as by location and type. However, ontologies serving diverse communities are often more complex than needed for specific applications, creating barriers to adoption by researchers and resource developers. For example, cell atlases often attempt simplifications by manually building term hierarchies linking to cell type and anatomy ontologies, but these may include relationship types unsuitable for grouping annotations.

We present tools for validating human expert curated term hierarchies, developed in two human reference atlas projects, against ontology structures. The tools provide tabular statistics plus graphical views of matching and non-matching terms and relationships to support discussion and conflict resolution. The HuBMAP Human Reference Atlas (HRA) effort is used to validate the approach and tools, and the Human Developmental Cell Atlas is featured as a use case.

## Introduction

### Ontologies and FAIR sharing

Biomedical ontologies are widely used to annotate data and analysis results to ensure they are Findable, Accessible, Interoperable and Reusable (FAIR)^1^. The meaning of ontology terms (the types of entity to which they refer) is made clear using referenced definitions, ensuring consistent use in annotation. The use of standard, resolvable ontology term identifiers in annotation—both within and between datasets—aids Interoperability. To support Findability, the FAIR principles require annotation with rich metadata (F2), indexed in a searchable resource (F4)^1^. Ontology-based annotation supports this by providing a rich set of standardized terms for annotation, including standard labels and curated synonyms. When these annotations are indexed in searchable resources, e.g., Chan Zuckerberg Cell by Gene (CZ CELLxGENE) Discover ^2^, they aid Findability by supporting search with a range of terminology found in the literature—for example, for cell types, cellular components, or tissues.

Ontology terms are embedded in complex graphs of relationships recording the relationships between the entities they represent. This ontology structure—logical relationships between terms using well-defined, standardized relations—allows researchers to take advantage of biologically meaningful groupings (e.g., grouping by anatomical location or cell type) to search, combine and analyze annotated data. For example, the Gene Ontology (GO) is widely used, along with a vast body of gene annotations, to detect enrichment of genes in experimentally derived gene lists by their molecular function, cellular location and the biological processes in which they participate^3^. This typically relies on graph representations of ontologies using the subset of the relationship types that relate specific classes to more general parent or ancestor classes (typically classification and parthood relationships). For example ‘enzyme activity’ sits above (classifies) ‘kinase activity’ in the graph; ‘immune response’ sits above ‘B-cell cytokine production’ and ‘mitochondrion’ site above mitochondrial parts such as ‘mitochondrial matrix’. In GO, all relationships are defined in the Relations Ontology^4^ which is shared with many other ontologies within the Open Biological and Biomedical Ontology (OBO) Foundry. The relationships represented in this graph are safe for grouping, not only because of the choice of relationships, but also because of their use in formal, logically quantified axioms. The part relationship between mitochondrial matrix and mitochondria records that *all* mitochondrial matrices are part of (some) mitochondrion. We can therefore conclude that if a protein is annotated as localised to the mitochondrial matrix, it is localised to mitochondria. If a relationship were used to indicate that some mitochondrial matrices were found in other structures, this conclusion would not be correct. In the OBO standard, this logical structure is defined using the W3C standard ontology language OWL^5^.

A similar approach is taken by resources using ontology structure to aid findability. For example, the CZ CELLxGENE Discover platform uses classification and part relationships in OBO standard ontologies to support structured search for datasets by cell type, anatomical structure, disease state, and anatomical stage. Other resources leverage OWL semantics directly for grouping, e.g., Virtual Fly Brain uses OWL queries to support user requests for 3D neuron images and connectomics data by neuron type, location, neurotransmitter and lineage^6^.

### Widely used ontologies inevitably become complex, so users need simplified views

Biomedical ontologies serve multiple research communities and disciplines with different needs and terminological preferences. Any well-used biomedical ontology also faces constant pressure to expand and keep pace with the growth of knowledge, to add new terms in response to the demands of curation and to support new communities and new applications many of which require different axes of classification. This results in increased complexity as well as increased size. For example, a neuron ontology supporting multiple communities of users may classify neurons by morphology (Martinotti), location (cortical layer 4), neurotransmitter (GABAergic), and marker expression (Somatostatin), electrophysiological properties (spiking) and transcriptomic similarity. Manual maintenance of multiple axes of classification is impractical in large ontologies, so sustainable ontology development requires the use of automated classification via the use of logical definitions and reasoning^7–10^. This typically requires the definition and use of more relationship types and formal axioms (property characteristics, hierarchies, and chains^5^), adding still more complexity and sometimes obscuring simple relationships. For example, the Uberon anatomy ontology uses a rich set of relationship types including many different part relationships (Fig. 1).

**Fig. 1.**
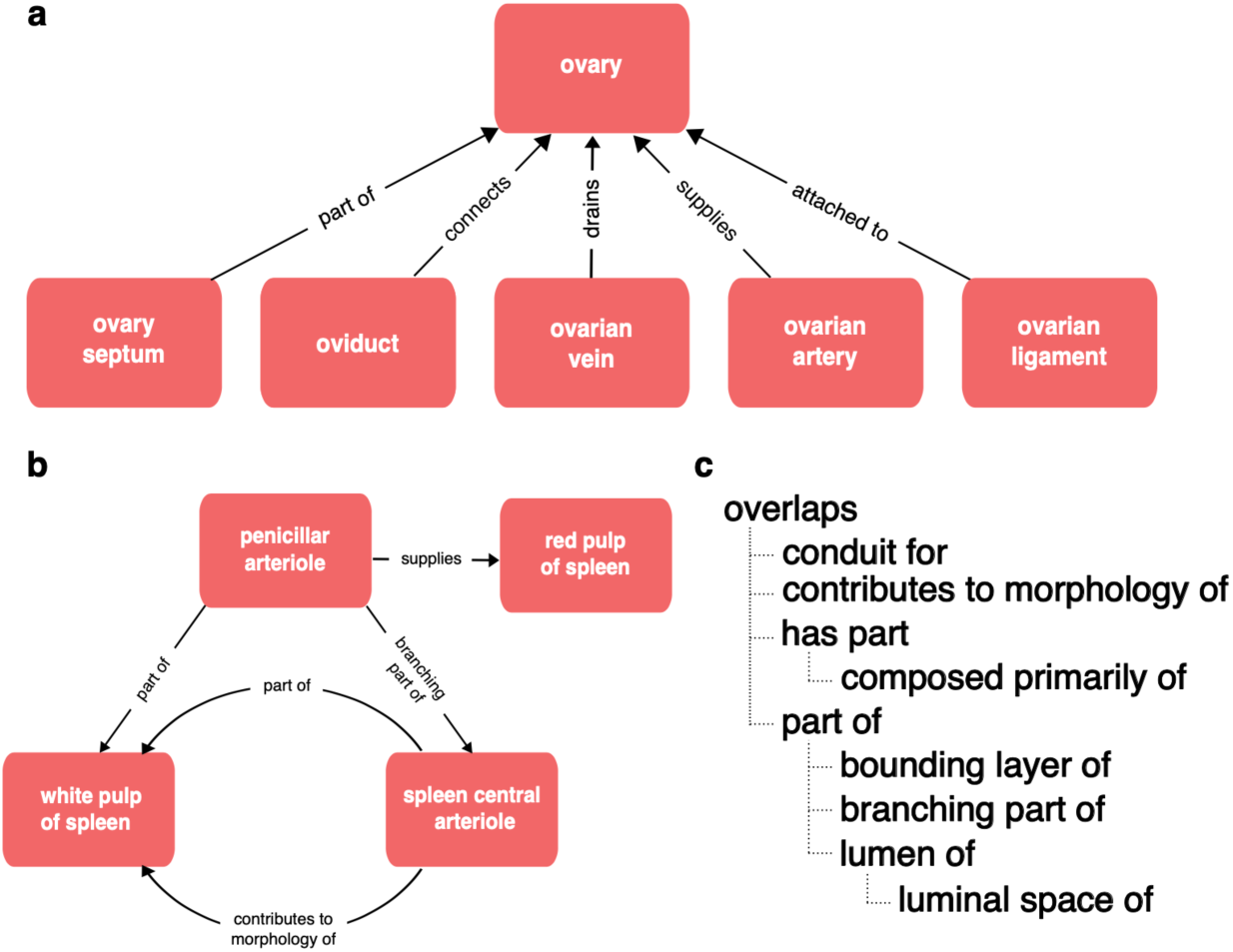
Relationships in Uberon. **(a)** The variety of relationships in Uberon, is illustrated using relationships between the ovary and various related anatomical structures, including relationships representing parthood, connectivity, attachment and supplying/draining vasculature. **(b)** The structure of relationships in Uberon is not limited to simple hierarchies (trees), illustrated here by the complex graph of relationships between spleen vasculature and the red and white pulp of the spleen. **(c)** Relations in Uberon are arranged in a hierarchy that can be used for inference, illustrated using the part relations hierarchy. If a lower relation in the hierarchy applies between two classes, then OWL reasoning infers a higher relation (e.g., and OWL reasoner would infer that the kidney capsule is part of the kidney from the relationship “‘kidney capsule’ bounding_layer_of some ‘kidney’” and the hierarchy “‘bounding_layer_of SubpropertyOf ‘part_of”).

This complexity is useful for driving automated classification, which is a key component of the pipelines used to ensure scalability of CL, Uberon, GO and many other OBO Ontologies^10,11^, adding thousands of inferred classifications. But not all relationships are useful for grouping annotations by anatomical location. A user querying for gene expression or cell types in the ovary (Fig 1A), will not expect to get results including gene expression or cell types in connected structures (ovarian ligament, oviduct, ovarian artery or vein). In the part relations hierarchy (Fig 1C), only part_of and its subproperties (e.g., bounding_layer_of) are completely safe for grouping annotations by anatomical location. For example, ‘finger’ part_of some ‘hand’ records that ‘all fingers are part of some hand’ so if a gene is expressed in a finger or some part_of a finger, it is safe to conclude that it is expressed in a hand. The ‘overlaps’ relationship records that two structures share some part, so if A overlaps B and B overlaps C, the part shared by A and B may not include the part shared by B and C. So it is not accurate to use overlaps for grouping annotations. For example, the ‘hepatic portal vein’ overlaps the liver (only part of it is in the liver), so using overlaps to group annotations to the liver could return annotations to regions of the vein outside of the liver.

This description applies to many highly-used ontologies including Uberon^12^, the Gene Ontology (GO)^13^, the Cell Ontology (CL)^10^, the Neuron Phenotype Ontology^14^, the Drosophila Anatomy Ontology^11^ and the Mondo Disease Ontology^15^.

While these ontologies are successfully used for annotation, search, and query by many different projects, they often have much more complexity than is strictly needed for individual use cases, which typically require only a subset of terms and relationships. In such cases, the full ontology, in all its complexity, can be intimidating for application developers to use. This is particularly an issue for projects that require a simple, minimal, browsable view of an ontology that fits their use case. We therefore need ways to generate simplified views of these ontologies in which all relationships are an accurate (if simplified) reflection of the content of the original ontology and that fit the use cases of the resource they are generated for.

### Informal annotation of atlases

Online anatomical and cell type atlases frequently make use of a hierarchical arrangement of terms used in annotation as a way to allow browsing of related content—sourcing terms and hierarchies from experts and/or basing them on data. Examples include the Allen Brain Atlas anatomical hierarchies used to annotate the standard anatomical atlases and parcellation schemes of the Allen Brain Atlas reference brains^16^; the Human Lung Cell Atlas—an atlas of transcriptomically defined lung cell types^17^; multiple transcriptomic cell type atlases developed for human, and mouse whole brains and brain regions^18,19^, hierarchies of Drosophila neuron types for annotation of connectomics^20^, the HuBMAP Human Reference Atlas (HRA)^21^, and the Human Developmental Cell Atlas (HDCA)^22^. Some hierarchies are derived from annotated reference data, e.g., representing transcriptomic similarity (Human Lung Cell Atlas, whole mouse brain), or anatomical part-whole relationships (Allen Brain Atlas) whereas others are based on expert opinion linked to publication and experimental evidence (HuBMAP HRA)^23,24^. These hierarchies are typically developed and shared in spreadsheets with relationships represented as duples—adjacency within a row specifies that two terms are in a hierarchical relationship of some kind—rather than triple with an additional slot for relationship type. However, additional Standard Operating Procedures (SOPs) or similar may specify a blanket interpretation of relationship type between adjacent terms. They are also almost always single inheritance (each term can have only one parent), which may benefit usability, but limits expressiveness.

The level of formalization varies greatly between these hierarchies—from completely informal with no mappings to ontologies (the current state of the Human Developmental Cell Atlas) to systems with extensive Standard Operating Procedures (SOPs), ontology mappings and some systems for specifying relationship types. For example, the Human BioMolecular Atlas Program (HuBMAP) Human Reference Atlas (HRA) uses a table-based system (ASCT+B tables) for building hierarchies of terms for anatomical structures (AS), cell types (CT) and Biomarkers (B)^21^. The terms used are mapped to Uberon^12^ (AS) or the Foundational Model of Anatomy (FMA)^25^ if an Uberon term does not exist, the Cell Ontology (CT) and HGNC^26^ (B). HRA SOPs name relationships that apply from AS to AS (ccf_part_of for organ tables, ccf_branching_part_of for vasculature tables), CT to AS (ccf_located_in), CT to CT (ccf_is_a) and CT to B (ccf_characterizes). While these are not formally defined or mapped to OBO relations, there is guidance for their use in SOPs^21,27^.

While these hierarchies are less formalized and simpler than those found in ontologies, they are a potential source of expert or data-driven content or correction for ontology structure. Conversely, the structure of ontologies can provide a means to validate the relations in these hierarchies against a range of curated, formally defined relationships and suggest missing relationships. This provides an important test for manually curated hierarchies built in spreadsheets with limited or no tooling support. Critically, it also provides the rigor needed to safely group annotations.

### Ubergraph

Ontologies encoded using standard formalizations such as OWL can be thought of as queryable classifications. Ontology terms refer to classes, are arranged in a classification hierarchy and are related to each other using standard relations (OWL objectProperties) and quantifiers. For example, ‘podocyte’ (CL:0000653) refers to the class of all podocyte cells, which are classified as epithelial cells. A relationship records that all podocytes are part_of some ‘glomerular visceral epithelium’ (UBERON:0005751), where part_of is a formally defined relationship in the OBO relations ontology^4^ and ‘some’ is a logical quantifier.

Not all relationships and classifications are stated directly. A chain of classifications in Uberon and CL connects ‘podocyte’ to ‘cell’, and a chain of part_of relationships connects ‘glomerular visceral epithelium’ to ‘kidney’. From this, we can infer that all podocytes are cells and are part of a kidney. Indirect relationships can also be inferred across relationship types based on hierarchies and rules that connect relation types. For example, the terms ‘kidney’ and ‘kidney capsule’ are related in Uberon using a specialized part_of relationship—bounding_layer_of (Fig. 1C)—from this we can infer that the kidney capsule is part of the kidney.

Ubergraph^28,29^ is a knowledge graph (triplestore) that integrates a large set of interconnected (mutually importing) OBO foundry ontologies, including the Cell Ontology and Uberon. It uses standard OWL reasoning software to directly assert indirect classifications and relationships in an easily queryable form, asserting, for example, podocyte subClassOf cell, podocyte part_of kidney, and ‘kidney capsule’ part_of ‘kidney’. In other words, Ubergraph takes the knowledge that is distributed throughout the set of large, complex and interconnected ontologies and associates it directly with each term. Ubergraph can be used to fulfill a number of the use cases discussed so far:

- Grouping annotations: Simple Ubergraph queries can, for example, find terms for all epithelial cells in the kidney. The output of this query can then be used as input to query a database of annotations with these terms. A REST API built on Ubergraph packages queries that support this and other use cases exists (http://grlc.io/api/INCAtools/ubergraph/sparql/#/default/get_cell_by_location).
- View generation: Ubergraph queries can be used to construct simple, minimal, browsable views of an ontology consisting of a limited set of terms and relationship types. By leveraging the inferred relationships in Ubergraph we can link these terms together into a graph even if they are not directly connected to each other in the ontology.
- Validation: Ubergraph can be used to test whether a set of relationships asserted in an outside resource are correct according to the ontologies in Ubergraph.

In this paper, we describe software libraries and pipelines that use Ubergraph to generate ontology views and validate user-generated hierarchies. We will illustrate their use with examples from the validation of HuBMAP ASCT+B tables and HDCA annotation term hierarchies. We also discuss how the resulting artifacts can be safely used for annotation grouping and the limitations of the current modeling of relationships in OBO anatomy and cell type ontologies.

## Results

### ASCT+B table validation

ASCT+B tables encode expert-curated relationships among anatomical structures, cell types, and biomarkers, which are essential for building the Human Reference Atlas and for driving findability of HRA data through HRA web interfaces and knowledge graph^30^. To test these relationships against well standardised, curated ontologies we built an automated analysis pipeline that uses Ubergraph to test whether one of a pre-agreed set of relation types applies for each curated relationship. We use the results to generate a simplified view of Uberon and CL to support Human Reference Atlas use cases, including supporting queries for HRA data via the HRA knowledge graph (add ref). This validation process is run weekly on more than 30 organ-specific ASCT+B tables, refined over four years, involving over 50 experts authoring and reviewing tables and over 100 users applying them for diverse atlas construction and analysis tasks^30^. Validation reports and term summaries are published to support ongoing refinement (https://hubmapconsortium.github.io/ccf-validation-tools/).

The pipeline reads in ASCT+B tables data using an HRA API (https://apps.humanatlas.io/asctb-api/), then uses a generic component to test the validity of relationships, expressed as simple “subject, relation, object” triples, reporting the results as tables and graphs. The code is available as a Python library (verificado, https://github.com/INCATools/verificado) and a templating system (validation-template, https://github.com/hubmapconsortium/validation-template) for configuring and running validation pipelines on GitHub.

The ASCT+B table format represents relationships between terms by their order in the row, with adjacent terms in a row forming an object (left) and subject (right) pair^27^. Fig. 2A exemplarily shows a small part of the Kidney table. The fourth data row records two relationships—one between the renal corpuscle and the nephron and another between the nephron and the kidney.

**Fig. 2.**
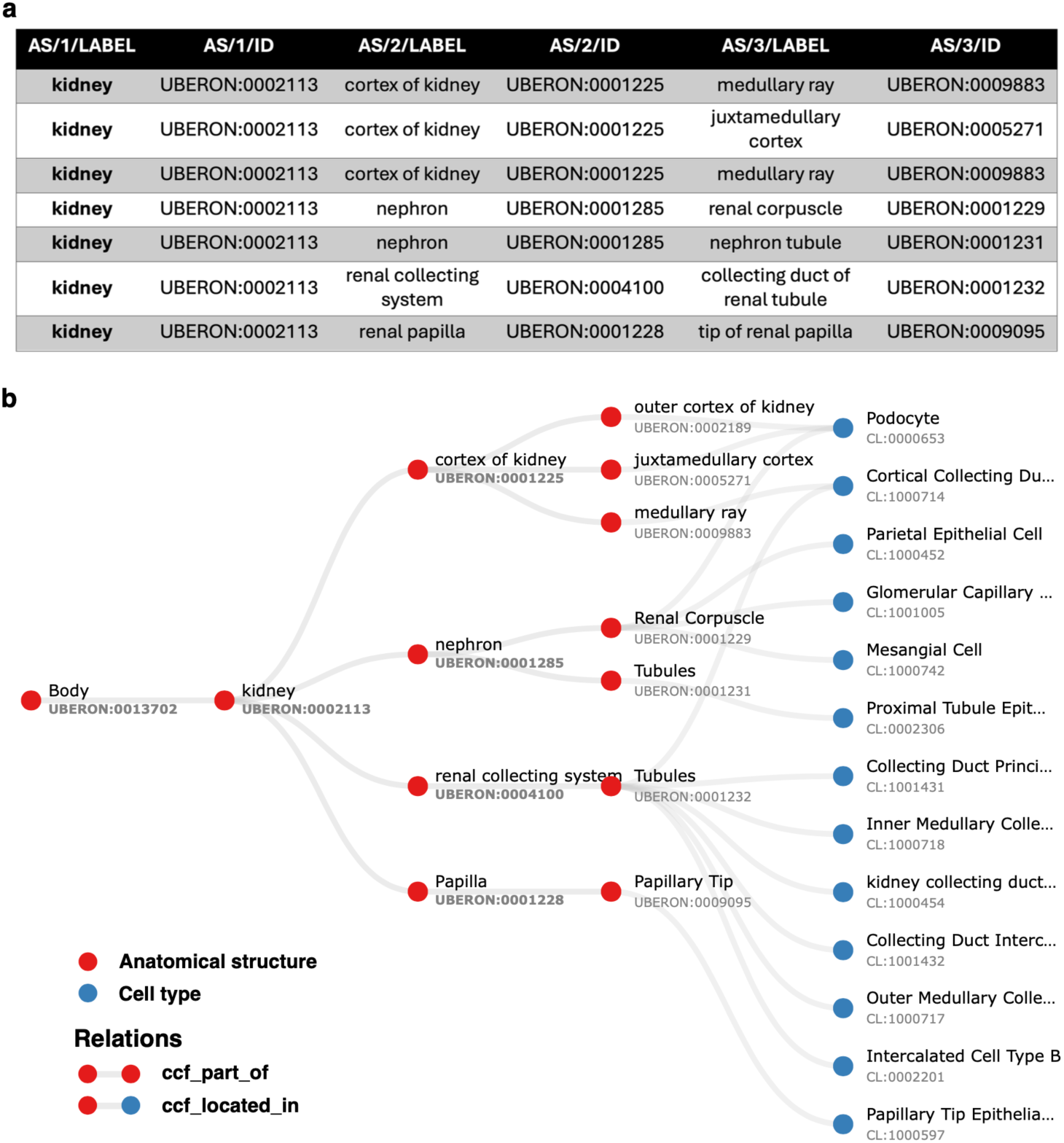
Kidney ASCT+B table relationships and graph representation. **(a)** A small part of the ASCT+B Kidney table (version 1.5^31^) recording relationships between kidney substructures. All of these correspond to part_of relationships in Uberon. In other cases, the corresponding relationship in Uberon is subClassOf, overlaps, has_part, or connected_to or does not validate. **(b)** Graph representation of a section of the ASCT+B Kidney table (version 1.5^31^) with anatomical structures denoted by red nodes and cell types by blue nodes. Labels in the graph are derived from the nomenclature chosen by the authors of the table. As a result, some structures, such as ‘Tubules’, may share the same name but are distinguished by different Uberon IDs to indicate their reference to distinct anatomical entities.

The pipeline tests whether relationships recorded in the tables between anatomical structures (AS) and cell types (CT) are true according to Uberon and CL by querying against Ubergraph (see Methods for details and all the relationships in Fig. 4 and Fig. S1 for examples). The tables themselves have standard operating procedures that detail the relations that should apply between all AS (ccf_part_of), between all CT (ccf_is_a), between AS and CT (ccf_located_in) and between B (Biomarkers) and CT (ccf_characterizes)^21^. These relations are bespoke to HRA (CCF refers to the HRA Common Coordinate Framework) and so do not match relations in the OBO relations ontology used by Uberon, CL, GO and many other OBO foundry ontologies^32^. On inspection, it became clear that mapping each of these four bespoke relations to a relation type used in Uberon and CL would not be sufficient for validation. For example, ASCT+B tables contain a mix of subClassOf and part_of relationships (e.g., Uberon models ‘glomerular visceral’ and ‘parietal epithelia’ are subclasses of ‘glomerular epithelium’, rather than parts of it). Occasionally, other relationships apply and are reflected in Uberon and CL as seen in the following examples: ‘ureter’ overlaps (has some part in the) ‘kidney’, ‘ovary ligament’ connected_to ‘ovary’; ‘myeloid cell’ develops_from ‘common myeloid progenitor’. The pipeline tests all of these, generating graphical reports of relationships that do not validate as shown in Fig. 3.

**Fig. 3.**
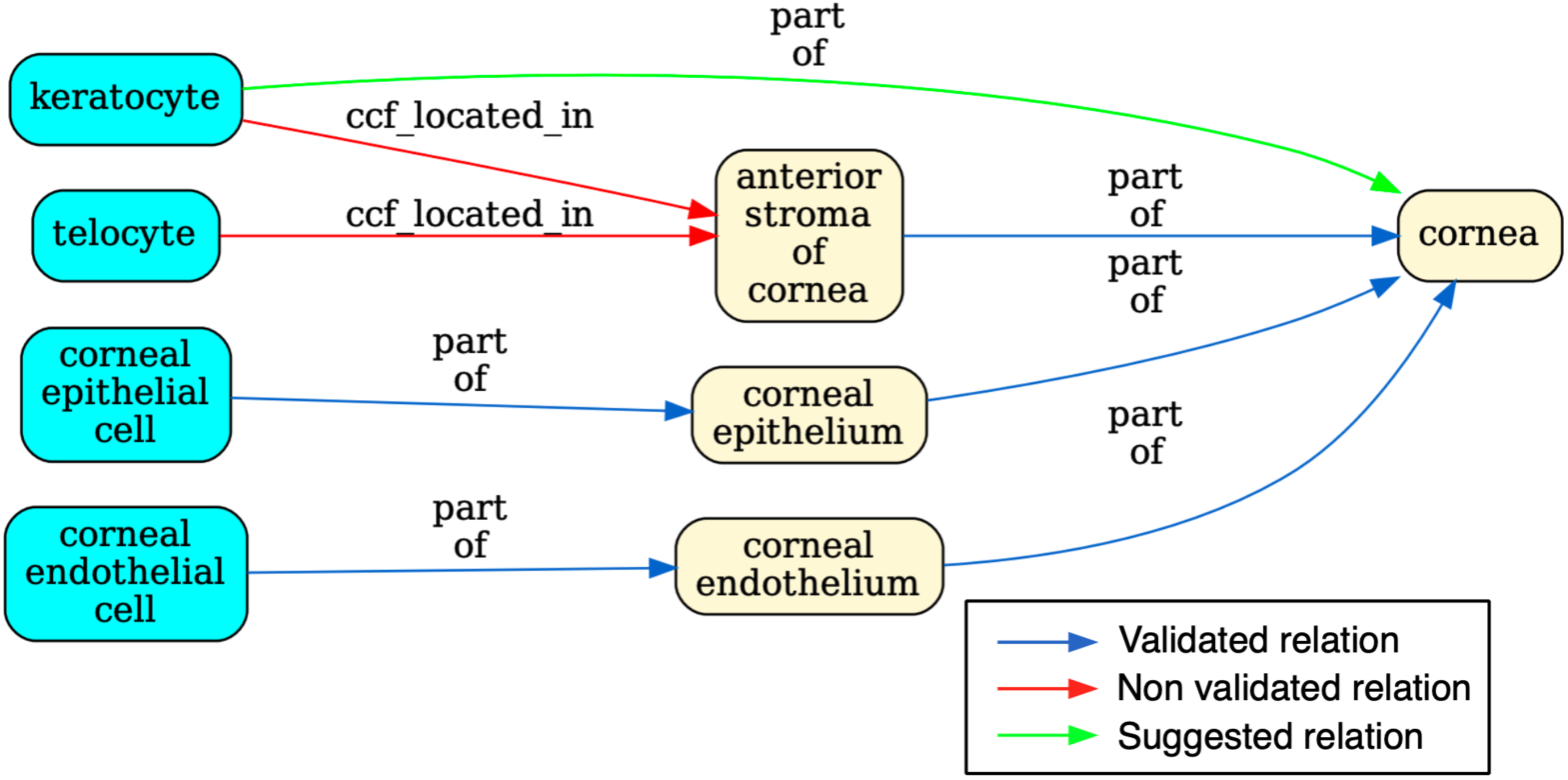
Validation for a portion of the ASCT+B eye table. The graph flags the relationship between ‘keratocyte’ and ‘anterior stroma’ of the ‘cornea’ in red to indicate that it is unsupported by CL/Uberon. Keratocytes are predominantly located in the anterior stroma, but not all are. Keratocyte is recorded in CL as part of the ‘substantia propria of the cornea’ which encompasses anterior and posterior stroma^33^. Because ‘substantia propria of the cornea’ is not used in the ASCT+B table, the validator suggests instead a part_of relationships to ‘cornea’ (in green), the most precise enclosing structure that is in the table. The authors of the table may choose to add this relationship or the term “substantia propria of the cornea” to their table in future, but without a change, this suggested (correct) relationship is added to the output ontology file. Telocyte relationship to “anterior stroma of the cornea” is also flagged. This is a more general cell type present in many other tissues, so a part_of relationship to “anterior stroma of the cornea” would not be correct (it would lead to incorrect grouping of annotations). However, as some telocytes are present in this tissue, it would be formally correct to add as has_part relationship between ‘anterior stroma of cornea’ and telocyte.

**Fig. 4.**
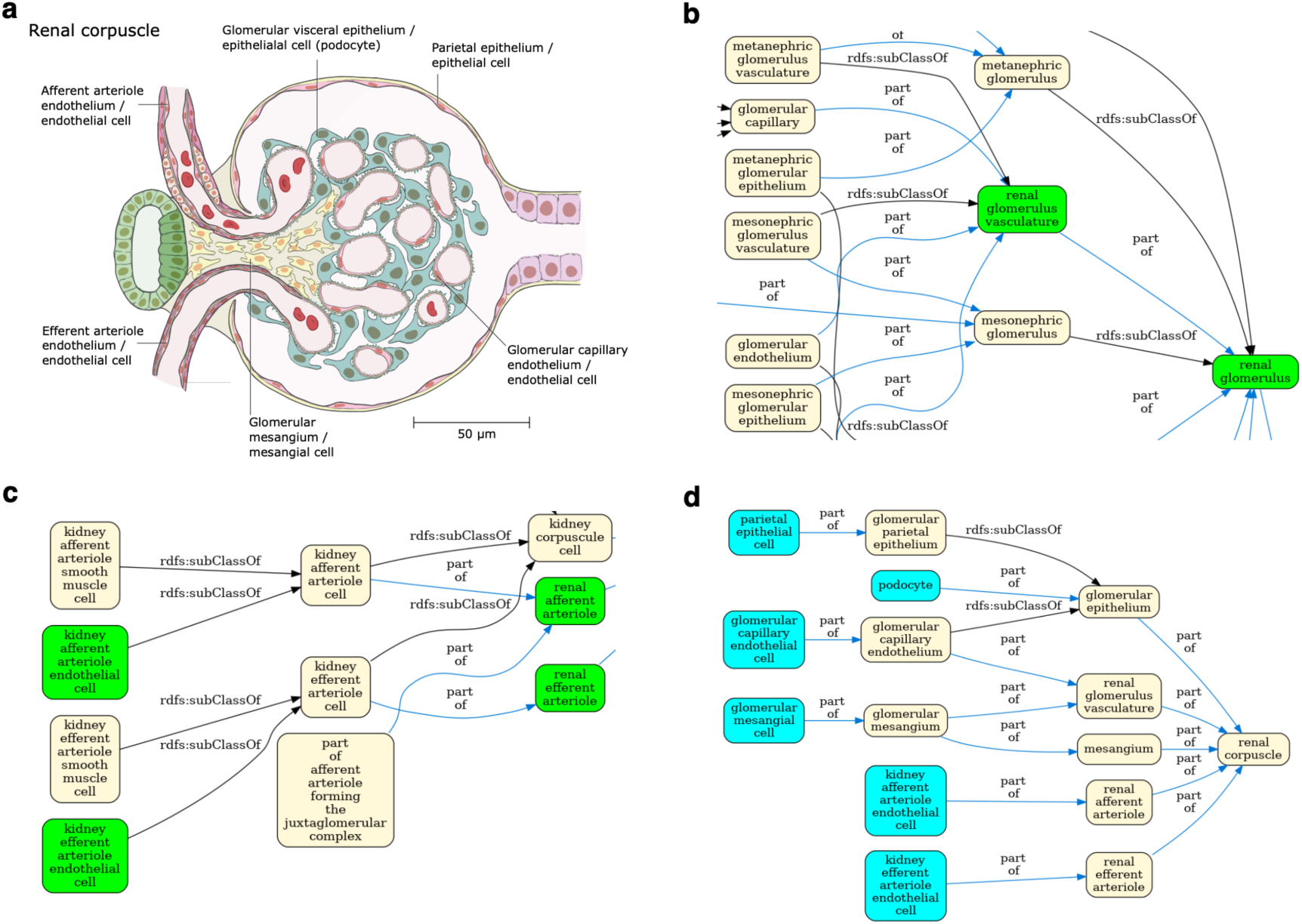
View generation for the renal corpuscle. **(a)** A 2D FTU illustration of a ‘renal corpuscle’ from the HRA^35^. **(b)** Enlarged detail of the Uberon/CL ontology graph for the renal corpuscle (complete graph in Fig S1). Terms referenced in the HRA kidney ASCT+B table in green. This detail features some of the many developmental terms that are out of scope for the HRA. **(c)** A different portion of the graph in Fig. S1 with grouping terms, such as ‘kidney afferent arteriole cell’, not needed by the HRA). **(d)** Final HRA view of the renal corpuscle with cell types in cyan and anatomy in yellow.

Table 1 shows another common pattern of non-validating relationships—a generic term under a specific one. The kidney ASCT+B table uses the term ‘endothelium’, which applies to endothelia all over the body, in between ‘kidney’ and the terms for various kidney endothelial cells such as the ‘vasa recta descending limb cell’. The authors clearly mean to refer to the specific endothelium of this structure, but this information is not accessible to the OWL model or simple graph reasoning. Expressed as a quantified statement in OWL (all endothelium is part_of some vasa recta descending limb) this is obviously incorrect. The use of this hierarchy for retrieving cell types in the kidney would return many endothelial cell types outside of the kidney.

**Table 1.**
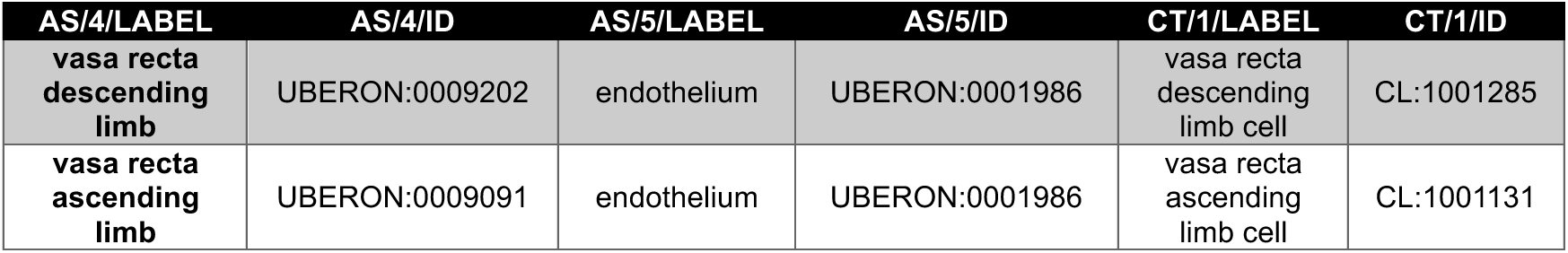
An extract of the Kidney ASCT+B Table (version 1.5^31^) containing the term ‘endothelium’. Although this term applies to all endothelia, in these two instances it is applied as the endothelium that is part_of the ‘vasa recta descending limb’ and part_of the ‘vasa recta ascending limb’.

| AS/4/LABEL | AS/4/ID | AS/5/LABEL | AS/5/ID | CT/1/LABEL | CT/1/ID |
| --- | --- | --- | --- | --- | --- |
| vasa recta descending limb | UBERON:0009202 | endothelium | UBERON:0001986 | vasa recta descending limb cell | CL:1001285 |
| vasa recta ascending limb | UBERON:0009091 | endothelium | UBERON:0001986 | vasa recta ascending limb cell | CL:1001131 |

Validated relationships are written to OWL. When validation fails, the pipeline finds the most specific valid relationship to another term in the table and writes this to OWL (see green arrow in Fig. 3 and description in legend). All original CCF relationships are also written to OWL as non-logical annotation axioms, preserving the original graph (Fig. 2B) for reference while avoiding potential incorrect logical inferences arising from non-validated relationships.

The result is a simplified view of Uberon and the Cell Ontology containing terms from the ASCT+B tables and relationships that are valid according to Uberon and CL. This simplification is illustrated in Figure 4 using the renal glomerulus (Panel A) as an example. Panels B-C illustrate the complexity of Uberon compared to the terms required by the HRA (highlighted in green). Panel D shows the simplified view of the Cell Ontology and Uberon generated using the HRA terms and relationships as input. Looking at the kidney more broadly, version 2.3 of the CCF ontology representation of the kidney has 124 terms, 84 relationships and 2 relation types, reduced from 236 terms linked by 384 relationships using 9 types of relation in Uberon and CL. The result is a much simpler ontology, removing terms that do not apply to adult *Homo sapiens*, such as the head kidney (UBERON:0007132), a part of the kidney in teleost fishes with an immune rather than renal function^34^, and terms referring to the anatomy of earlier stages of kidney development, such as the mesonephric renal vesicle (UBERON:0005331) and pronephric duct (UBERON:0003060).

Non-validated relationships are investigated as potential edits to Uberon, CL or the ASCT+B tables. In some cases, lack of validation points to a simple error in the original table, e.g., a syntactic or typographic error in an identifier that can then be corrected by ASCT+B table authors in future HRA releases. In other cases, it points to a dispute of biology or its ontological modeling that needs discussion to fix, or to a missing relationship in CL or Uberon that needs adding. For example, in version 1.0 of the eye table^36^, the authors included a relationship between the ‘neuron projection bundle connecting eye with brain’ (UBERON:0004904) and the sclera (UBERON:0001773). However, while this bundle passes through the sclera, it is not part of it. Based on guidance from CL editors, the authors removed the relationship in subsequent versions of the table to reflect a more accurate anatomical and ontological interpretation. Monthly HRA working group meetings are used to resolve disputes and misalignments; visual representations of the anatomical structure partonomy and cell type typology are critical for conflict and error identification and correction.

The result is an ontology with a mix of validated relationships using well-defined OBO relations. Relationships using a subset of these relation types can group anatomical structures, cell types, and data annotated with them by location. Other relationship types represented in the ontology could support other anatomical query use cases, for example, queries for connected structures.

### Improving the Human Developmental Cell Atlas

The Human Developmental Cell Atlas (HDCA)^22^ effort manually developed an anatomical structure-cell types-molecules hierarchy which contains a mix of relationship types, including connectivity (e.g., nerves to brain regions), subclass, overlap, part_of, adjacency, and developmental relationships. Applying the validation tools to this hierarchy illustrates how ontology-based validation helps clarify which relationships can reliably support grouping annotations by anatomical location, and where refinements to the hierarchy or the reference ontologies are needed.

For example, the HDCA hierarchy has the midbrain-hindbrain boundary under hindbrain. However, the isthmus organizer resides within the boundary, and signaling within this region organizes development of the adjacent parts of the human brain^37^. The isthmic organizer straddles the midbrain-hindbrain boundary and so has an overlapping relationship to hindbrain in Uberon (Fig. 5B, 5C).

**Fig. 5.**
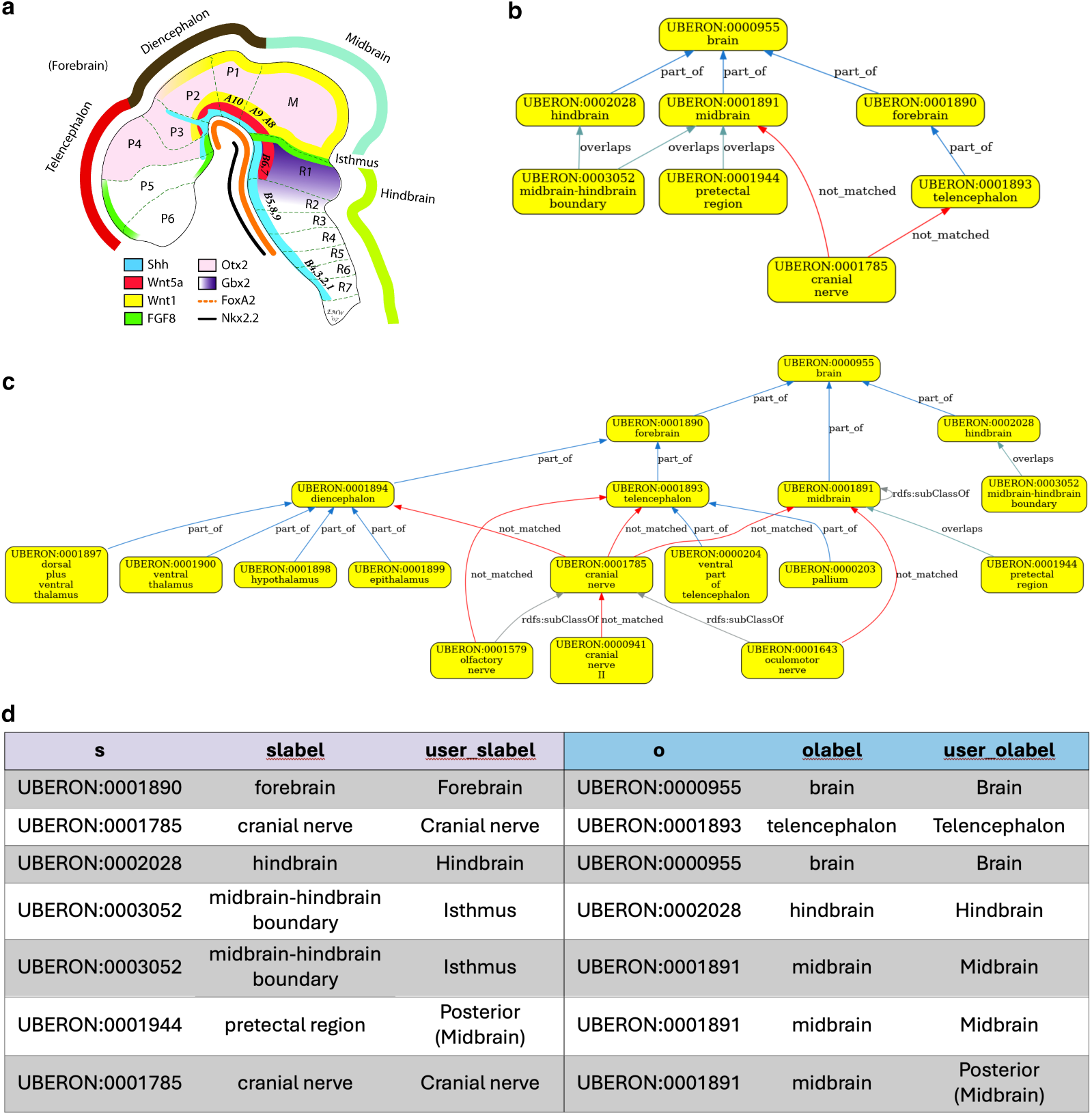
Validation of the midbrain-hindbrain bordering for HDCA. **(a)** Schematic of the developing brain. Note the Isthmus, midbrain-hindbrain boundary and the different expression of key developmental cell-cell signaling molecules on either side. Figure is adapted from Wexler and Gaeschwind, 2007^38^. Uberon reflects this by having *overlaps* relationships between the boundary and both midbrain and hindbrain (Panel b). **(b and c)** Validation graph for HDCA. Note that the nerve relationships (connectivity) do not validate because Uberon records these more precisely. Validation could be achieved by extending Uberon with a rule that infers connection to a whole from connection to its parts. **(d)** Table of asserted relationships in the HDCA hierarchy without relationship type specified.

The boundary forms in the neural tube and patterns the adjacent midbrain and hindbrain through the coordinated expression of key transcription factors and signaling molecules like Fgf8 (in the hindbrain side of the boundary) and Wnt1 (in the midbrain side of the boundary) and is a compartment boundary (cells do not cross it during development) (Fig. 5A).

If queries for cells in the hindbrain return cells at the midbrain-hindbrain boundary, this will include cells from the part of the boundary that is in the midbrain (e.g., the Wnt1 expressing cells on the midbrain side).

In some cases, developmental relationships apply. For example, HDCA has terms for ‘pharyngeal arches’ under ‘neural crest’. Neural crest cells are pluripotent cells that arise from the dorsal neural tube and contribute to many structures including the pharyngeal arches, which also contain cells originating outside the neural crest^39,40^. Uberon includes a ‘has developmental contribution from’ between ‘neural crest’ and ‘pharyngeal arch’ (Fig. 6). This developmental relationship is not suitable for grouping annotations as it would group annotations to neural crest with those to cell types that do not originate in the neural crest. In this particular case, if the authors of the table preferred a hierarchy based solely on grouping classes, an ontology editor could advise HDCA to use the grouping term ‘structure with developmental contribution from neural crest’ which is a parent class of ‘pharyngeal arch’ and is suitable as a general term for annotation and grouping by type.

**Fig. 6.**
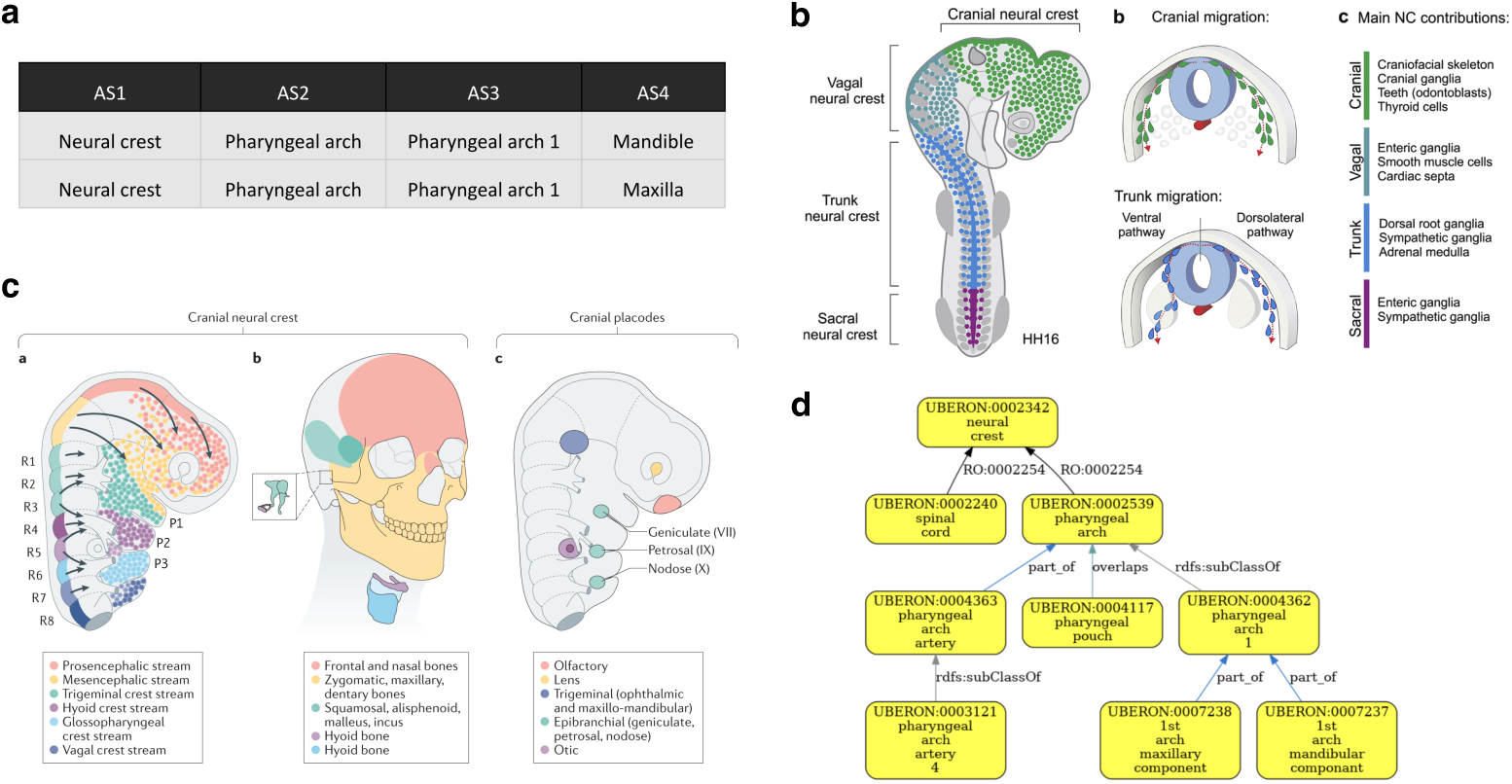
Validation of developmental contributions of neural crest to pharyngeal arch structures. **(a)** HDCA hierarchy with terms for pharyngeal arches under ‘neural crest’. AS: Anatomical Structure. **(b)** The neural crest arises from the dorsal neural tube and migrates to contribute to many tissues throughout the body, not just the pharyngeal arches. Figure is adapted from Rohstein et al., 2018^39^. **(c)** Neural crest migration in the head, including migration to pharyngeal arch 1, which forms structures including the upper and lower jaws (maxilla and mandible in HDCA hierarchy) and the bones of the middle ear. Figure is adapted from Martik and Bronner, 2021^40^ **(d)** Validated Uberon hierarchy showing the relationship has_developmental_contribution_from (RO:0002254) between ‘pharyngeal arch’ and ‘neural crest’, illustrating that HDCA hierarchy includes developmental relationship without necessarily including higher-level grouping terms.

**Fig. 7.**
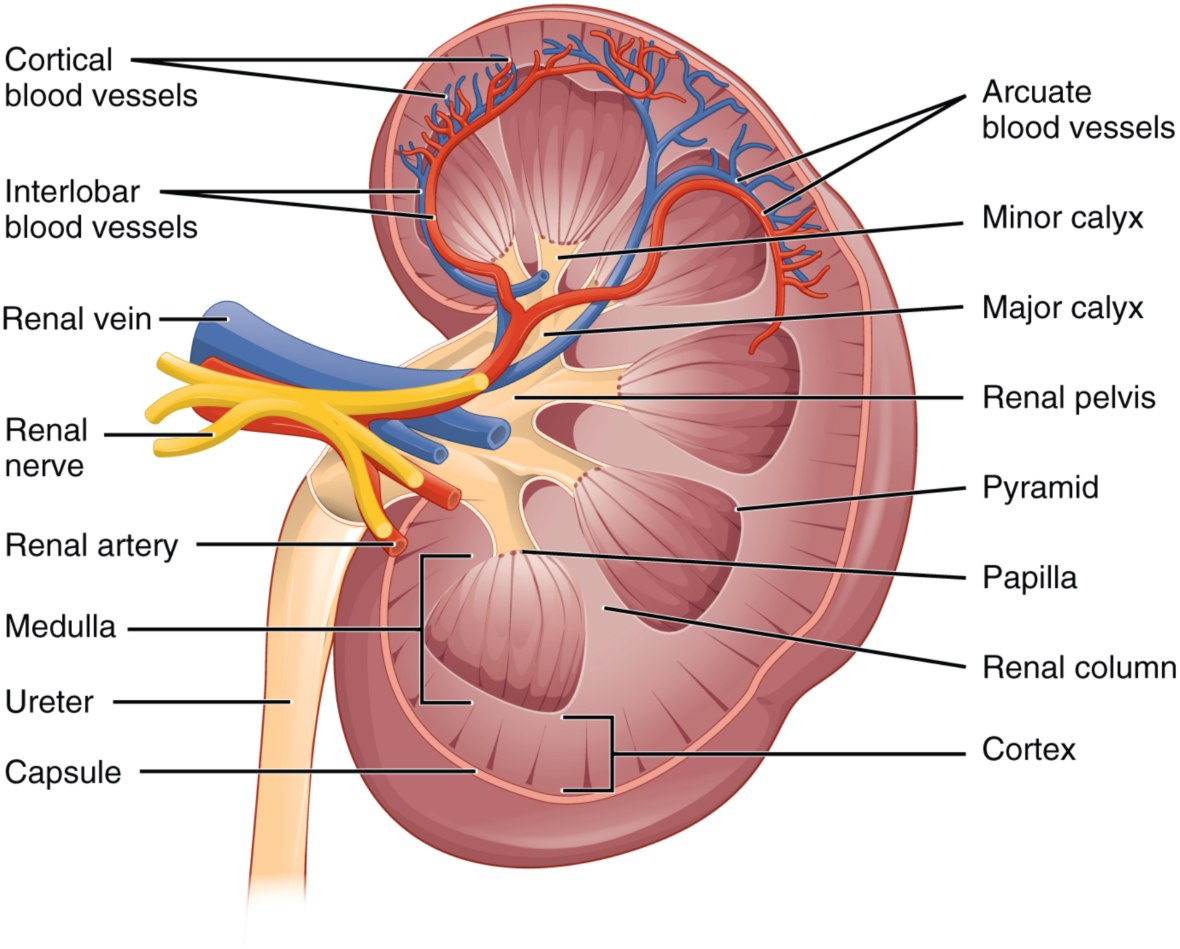
Schematic of coronal section of the kidney^42^. As the renal columns have the same internal structure to the cortex, some treatments consider them to be part of the cortex while others (including those who annotated this figure) do not.

As well as improvements to the HDCA, this work also identified issues with Uberon, such as the representation of the thalamus. Although Uberon includes terms such as ‘dorsal thalamus’, ‘ventral thalamus’, and ‘dorsal plus ventral thalamus’, which users are already using to annotate thalamic structures, it lacks a term for ‘thalamus’ itself.

### Community tools

The problems and solutions we describe for working with the HuBMAP HRA and HDCA are similar to those we have also encountered with other projects. For example, groups working on annotation of integrated datasets for the Human Cell Atlas, on *Drosophila* cell types and mammalian brain region have all generated their own hierarchies for use cases where ontology hierarchies already exist. We have developed a suite of Python tools to make it easy for the wider community of ontology users to take advantage of ontology structure to validate their own hierarchies or seed ontology views.

verificado (https://github.com/INCATools/verificado) takes a set of relations and a configuration file specifying relation types to check. It reports on which relationships are valid, according to Ubergraph. The set of relations can be tailored to use cases, for example a user might choose to limit relations to those useful grouping annotations.

ubergraph2asct (https://github.com/hubmapconsortium/ubergraph2asct) takes a list of terms and a list of properties and generates reports of ontology (view) structure in a simple table format based on the HuBMAP ASCT+B tables. This facilitates an easy review of implicit ontology structure.

validation-template (https://github.com/hubmapconsortium/validation-template) is a template for creating a GitHub repository with a pipeline that generates graphical and table-based validation reports similar to those generated by ccf-validation-tools. Generation is driven by GitHub actions or can be run locally using a docker container that wraps all dependencies.

We also packaged the view-generation strategy described in this paper into the popular ROBOT ontology workflow tool^41^—where it is available via the ‘subset’ option in the ‘extract’ command.

All of the tools described here rely on Ubergraph, as it is the only knowledge graph that provides a unified, reasoned view of all OBO foundry ontologies and their interrelations, enabling consistent validation and querying of ontology-based structures.

## Discussion

The work described here demonstrates an effective method for the collaborative development and refinement of ontologies using existing hierarchies of user-defined annotation terms used in atlasing. In addition to enabling ontology extension, this approach also supports iterative improvement of the hierarchies themselves, allowing experts to revise and validate relationships. Validation tools make it possible to flag possible errors in expert authored hierarchies or missing terms in existing ontologies. By making the semantics of relationships in these hierarchies clear, our approach supports effective and accurate grouping of annotations by location and type while preserving other types of relationships, e.g., developmental relationships or anatomical connectivity.

### Challenges for validation

#### Modeling differences arising from terminological variation

There will always be competing terminologies and differences in terminology usage in anatomy and medicine. Ontologies generally deal with this by attempting to choose the dominant usage/terminology and capturing alternatives as synonyms with attached scopes: exact; broader; narrower; and related. When two separate groups of experts make decisions about how to structure an ontology, it is inevitable that they will sometimes make different, but equally justifiable choices about precisely what commonly used terms refer to. For example, all standard atlases of the kidney divide it into cortex and medulla, but do not all agree on where to draw the boundary between the two. In some treatments, the cortex refers to the entire outer region of the kidney, and the medulla is the entire inner region. The inner region includes columnar structures (renal columns) whose internal structure matches that of the cortex. Because of this, some treatments include the columns as part of the cortex, not the medulla, despite their inner location. Uberon follows this treatment, while the ASCT+B Kidney table follows the former. As a compromise, we added two new terms to Uberon, ‘cortical region of kidney’ and ‘medullary region of kidney’ to map to the ASCT+B tables, supporting both use cases.

#### The relationship of cell types to anatomical structures

One important use case for ASCT+B tables (and many atlases) is to support listing of cell types present in specific anatomical structures. The reference ontology used, Uberon, represents this as part_of relationships where all cells of the referenced type are part of the referenced anatomical structure (e.g., all podocytes are part of a renal glomerulus) but otherwise uses has_part. However, Uberon includes relatively few *has_part* links to cell types. To address this gap, ASCT+B tables have provided a valuable source of expert input, which we have used to add new *has_part* relationships to Uberon. For example, based on the lung ASCT+B table v1.3^43^, we added 22 *has_part* assertions (https://github.com/obophenotype/uberon/issues/2960). This reflects cases where the same cell types occur across structurally repeated segments of the lung. Rather than creating redundant cell-type terms for each segment, we modeled this more efficiently by asserting that these lung structures has_part each relevant cell type.

ASCT+B tables collect markers attached to cell types in the context of specific tissues. It is not always clear whether the markers collected are tissue-specific or not, but many are. In some cases, organ/tissue experts intentionally provide markers that encompass a biomarker set which uses both general and tissue-specific markers to define cell types. For instance, general markers are used to define epithelial tissue/cells, but additional markers are added to provide specificity to which type of epithelial cell. In the lung table v1.5^44^, for example, the authors list general epithelial markers for club cells, such as CDH1 and EPCAM, alongside more specific markers like SCGB1A1 and SCGB3A2. Using combinations of these markers plus the spatial context in the tissue allows researchers to accurately identify cell types. Anatomical context is especially important when linking markers to cell types because most cell types are highly specialized and organ-specific. Recent analyses across large-scale single-cell datasets, such as Tabula Sapiens, reveal that approximately 64% of cell types are unique to a single organ^45^, highlighting the need to consider tissue environment when assigning biomarkers. While some cell types and markers are shared across organs, the dominant pattern is one of organ-specific specialization. This underscores the importance of allowing organ experts to combine general and tissue-specific markers in light of the anatomical context to achieve biologically meaningful classifications.

One solution to this would be to generate tissue-specific cell-type terms for every instance of a generic cell type linked to a specific tissue in the tables. However, this would massively bloat the ontologies—making manual maintenance of said ontologies more difficult—and some of these compound terms can end up with very long, unwieldy names, for example, ‘kidney loop of Henle long descending thin limb outer medulla epithelial cell’. Another approach is to incorporate information about the cell type location into the axiomatization of marker sets. This latter approach is presently being adopted by the HRA for representation in OWL and integration into the HRA Knowledge Graph. Specifically, instances capturing the study discovery of the marker set are created, with each instance giving pointers to the targeted cell type, its tissue location, the defining markers and supporting references that substantiate these findings^30^.

#### Resident immune cells

All human tissues have immune cells. In every organ, except the central nervous system, the immune cells present include non-resident as well as resident immune cells, with resident cells typically having distinct phenotypes and expression profiles^46^. While there are rapid advances in our understanding of resident immune cells and our ability to identify them, this is an active field of research. As a result, distinguishing resident from non-resident immune cells in individual tissue samples is challenging.

This, in turn, poses a challenge for building ontologies of anatomy and cell types to be used for annotating the cell types found in tissue samples. part_of relationships record that *all* cells of some specified type are part of a particular type of anatomical structure (all podocytes are part of some glomerular epithelium). Creating subclasses of general immune cell classes for every possible location would bloat ontologies enormously, and the classes created would cover both resident and non-resident immune cell types in those locations. One way to get around this is to add has_part relationships between anatomical structures and cell types (e.g., recording that the ‘kidney interstitium’ has_part some (types of) ‘B cell’).

ASCT+B tables—per SOP instructions—aim to exclusively record resident (immune) cells where this distinction can be clearly made. When comparing ASCT+B tables with experimental data, transient (non-resident immune) cells need to be excluded to arrive at a perfect match.

Some resolution of this issue is likely to come from increased knowledge of the markers characteristic of resident immune cells, allowing us to add specific cell types for these along with clear criteria for identifying them.

### Advantages and limitations of table-based approaches to building hierarchies

Simple hierarchies (single inheritance tree structures) are a simple and intuitive way to structure information. They are much easier to browse and visualize than multi-inheritance graphs with many relationship types. Spreadsheets and other tables (CSVs, DataFrames) are the default medium for most biologists’ work. It is therefore not surprising that spreadsheet-based representations of simple hierarchies are a common starting point for biologists aiming to organize annotation terms into a form that is easily browsable and visualisable.

This approach, combined with SOPs to guide and constrain the work of expert contributors and visualizations of resulting graphs with multiple node types and diverse edge type using the ASCT+B Reporter^23^ has enabled dozens of organ experts across multiple consortia to collaborate and rapidly expand the information in the ASCT+B tables.

Even with SOPs, any system that relies on manual curation into spreadsheets by a diverse set of expert contributors will accumulate errors and inconsistencies. Validation against ontologies provides a means to find and correct these and provide feedback to ontologies on missing or incorrect relationships.

A major limitation of this approach is that it fails to represent multiple relationship types between similar types of entities or to represent multi-inheritance hierarchies even where those may be useful. Biologically relevant ways of dividing up a structure may not conform to single inheritance. For example, the intestine can be divided into layers (e.g., mucosa, submucosa, muscularis) and segments (e.g., ileum, jejunum). A structure such as the submucosa of ileum belongs simultaneously to both: it is part of the ileum (segment) and a subclass of submucosa (layer). This dual inheritance cannot be cleanly represented in a single-parent spreadsheet format. Similar issues arise in cell type classification (where cells may be classified by location, lineage, and function). These complexities are better represented using ontology-based graphs that support multiple inheritance and diverse relationship types.

An alternative approach would be to start from a set of terms needed for annotation, selected by experts and to generate ontology views from these (which can be multi-inheritance), along with visualizations and draft tables. For view generation, we have developed tooling for the ROBOT ontology manipulation tool^41^ that allows the generation of minimal ontology views using a set of terms and relationships as input. The resulting view can then be visualized as a graph and used to generate tables (inspired by the ASCT+B table structure) via the ubergraph2asct (https://pypi.org/project/ubergraph2asct/) package, for review by experts who can suggest missing relationships and point out incorrect ones. With the existing ontology structure as a starting point, this approach is potentially a more rapid way to reach consensus while supporting interaction via spreadsheets.

This approach addresses the limitations of table-based hierarchies by leveraging the ontology’s existing multi-inheritance structure and logical relationships. It avoids ontology bloat by representing spatial context rather than creating numerous compound classes. The resulting ontology views preserve the expressivity needed to capture anatomical and functional complexity (e.g., intestinal layers vs. segments, or immune cell residency) while remaining accessible through table-based visualizations. This supports expert review and iterative refinement, facilitating both validation and knowledge integration in a more scalable and semantically robust way.

#### A toolkit for constructing biomedical atlases

The development of biomedical atlases is becoming a pillar of modern medical and life sciences research, where they are invaluable resources for providing access to data to help understand complex biomedical problems. However, the process of building these atlases often faces challenges in terms of data standardization and interoperability. The Human Cell Atlas project has emphasized the need for standardized cell type annotations^47^. The ontology validation tools that we present here simplify the complexity by allowing researchers to validate their simplified hierarchies against comprehensive ontologies. By improving accessibility and accuracy in ontology usage, this toolkit has the potential to accelerate research across multiple fields of biology, from developmental studies to disease research. As atlas projects continue to grow in scope and complexity, using the tools described here will be essential in maintaining data quality and interoperability, ultimately leading to more robust and impactful scientific discoveries.

Beyond validation, this study also demonstrates that, given appropriate tooling, informally structured hierarchies can be an effective mechanism for collecting knowledge from subject matter experts (SMEs). This approach is helpful for specifying the parts of an ontology required for a particular use case and as a source of information for improving ontologies. The graphical reports generated are valuable for discussion with SMEs to reconcile differences in representation. Additionally, the ontology views produced by our tools and pipelines provide a product that can accurately group content while being tailored to the needs of individual projects.

## Methods

The Ubergraph redundant graph^28^ is a comprehensive precomputed graph that explicitly includes all asserted and OWL-inferred existential restrictions, represented as simple triples combined with a complete transitive closure of the subClass graph. These inferences arise from property chains, transitive properties, property hierarchies, and other OWL axioms, and include relationships that are asserted rather than explicitly stated in the original ontologies. For example, inferred relationships include those inferred from chains of transitive relationships (e.g., if A part_of some B and B part_of some C then A part_of some C), classification (e.g., if A part_of some B and B subClassOf C then A part_of some C) and property hierarchy (e.g., if A bounding_layer_of some B and bounding_layer_of is a subProperty of part_of, then A part_of some B).

We use SPARQL queries of the Ubergraph redundant graph to test whether assertions about the relationships between terms found in ASCT+B are true (valid) according to the current structure of Uberon and CL ontologies. True relationships are added as existential restrictions to the Human Reference Atlas Common Coordinate Framework Ontology (CCF.owl) application ontology^48^. These axioms are annotated to record validation status and date, while non-validated relationships are used to generate reports for further investigation. In order to avoid producing a view of Uberon and OWL with terms lacking any relationship to other terms, we find the most specific valid relationship to another term in the graph for every relationship that doesn’t validate. All relationships in the ASCT+B tables are also written to the output ontology as simple triples using official ASCT+B table relations as OWL Annotation Properties. This ensures that the graph structure encoded in the ASCT+B tables is preserved in a form that is outside of the logical axiomatisation of the ontology, and so does not interfere with standard logic-based queries of the ontology.

We access a digested version of these tables via the ASCT+B API (https://apps.humanatlas.io/asctb-api/). For each pair of anatomical structure terms, we test whether valid subClassOf, part_of (BFO:0000050), overlaps (RO:0002131) or connected_to (RO:0002170) relationships exist. For each cell type (CT) anatomical structure (AS) pair, we test whether part_of or overlaps applies between CT and AS and if not, whether has_part (BFO:0000051) applies between AS and CT. Relationships between cell type terms are tested for subClassOf or develops_from (RO:0002202).

Ontology editors triage and turn the validation output files into reports (https://hubmapconsortium.github.io/ccf-validation-tools/), advising on the creation of new terms and relationships or corrections to tables as needed.

CCF-validation-tools is written in Python and uses Makefiles and Robot^41^ to run the validation and build the pipeline. Graphs are produced using the og2dot javascript library (https://github.com/INCATools/obographviz) combined with dot format to transform the graph into PNG and PDF. The pipeline is run weekly under the control of GitHub actions, automatically importing the latest ASCT+B table versions.

### Example of SPARQL queries for relationship validation

To test whether candidate relationships from the ASCT+B tables are valid according to the current structure of Uberon and CL ontologies, we run SPARQL queries on the Ubergraph redundant graph. For each relationship type (e.g., part_of, overlaps, connected_to), we query sequentially over all candidate subject–object pairs.

If a pair is returned in the result, this indicates that the triple <subject relationship object> is present in the ontologies and is therefore considered valid. If it is not returned, the next relationship type in the validation order is tested until all options are exhausted.

For example, the SPARQL query to validate a part_of relationship is:

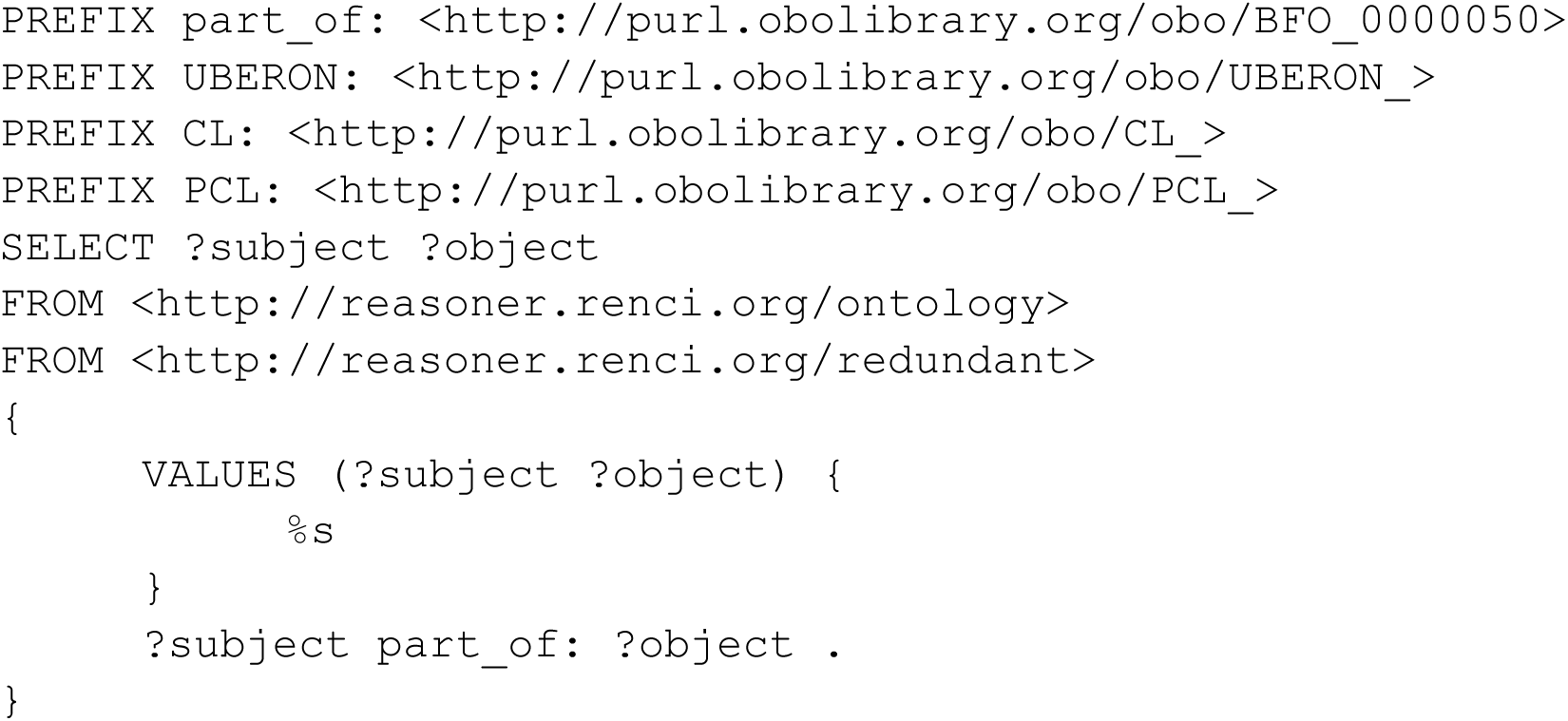

During validation, %s is replaced with the actual subject–object pairs extracted from the ASCT+B tables.

All SPARQL queries used in the validation process are available at: https://github.com/hubmapconsortium/ccf-validation-tools/blob/90e40ed0c82d2fb86735b48e3638803ecfd571af/src/uberongraph_tools.py#L9-L298.

## Data Availability

The ASCT+B validation pipelines described here are a major component of the Human Reference Atlas Common Coordinate Framework Ontology (CCF.owl) application ontology^48^. The examples in this paper correspond to version 2.3.0, available from https://bioportal.bioontology.org/ontologies/CCF.

All ASCT+B tables can be downloaded from https://lod.humanatlas.io/asct-b/.

## Code Availability

ubergraph2asct: https://github.com/hubmapconsortium/ubergraph2asct

Python library that queries Ubergraph with a list of terms and generates a simplified version of the ASCT+B table. License Apache 2.0. Available in PyPI https://pypi.org/project/ubergraph2asct/.

verificado: https://github.com/INCATools/verificado

Python library that verifies the relationships between pairs of terms against the ontologies available in Ubergraph. License Apache 2.0. Available in PyPI https://pypi.org/project/verificado/.

validation-template: https://github.com/hubmapconsortium/validation-template Github template to quickly validate an ASCT+B table and generate a graph view.

asct-parser: https://github.com/hubmapconsortium/asct-parser

Python library that converts an ASCT+B table into a TSV table with the format needed for the verificado library. License Apache 2.0. Available in PyPI https://pypi.org/project/asct-parser/.

All code for validating ASCT+B tables can be found at ccf-validation-tools: https://github.com/hubmapconsortium/ccf-validation-tools, published under an Apache License 2.0 license.

## Acknowledgements

This work was funded by NIH via the HuBMAP project (OT2OD033756 and OT2OD026671), the Cellular Senescence Network (SenNet) Consortium through the Consortium Organization and Data Coordinating Center (CODCC) under award number U24CA268108, the NIDDK under awards U24DK135157 and U01DK133090, and a Kidney Precision Medicine Project award U2CDK114886. JB’s work was funded by a gift from Schmidt Futures (https://www.schmidtfutures.org) to support the ‘Cell Annotation Platform’. ARC, APB, HK, PR, HP, JAM, and DOS were supported by EMBL-EBI core funds.

## Author contributions

***Anita R. Caron*** wrote the majority of the software described in this paper and contributed to writing.

***Aleix Puig-Barbe*** was responsible for much of the work on the Cell Ontology and Uberon arising from the use of the validation pipeline described in this paper. He also contributed significantly to writing and constructed the figures in this paper.

***Ellen M. Quardokus*** carried out the majority of the work on ASCT+B table construction with subject matter experts (SMEs), submitted new term requests to ontologies and correction and gave invaluable feedback to Cell Ontology editors. She also contributed significantly to writing the paper.

***Josef Hardi*** developed downstream software that utilizes the output of this work to construct the Human Reference Atlas Knowledge Graph (HRA KG).

***James P. Balhoff*** developed and maintains Ubergraph and the reasoning software used to create it.

***Jasmine Belfiore*** contributed to writing and editing this paper and ran the HDCA hierarchy validations.

***Nana-Jane Chipampe*** provided HDCA examples and biological expertise.

***Bruce W. Herr II*** developed and implemented the HRA Knowledge Graph compilation, enrichment, and publication process.

***Huseyin Kir*** contributed to software development.

***Mark A. Musen*** contributed to writing and editing the manuscript

***Paola Roncaglia*** was responsible for initial work on the Cell Ontology and Uberon arising from the use of the validation pipeline described in this paper, esp. re. thymus and lymph nodes.

**Helen Parkinson** contributed to writing the manuscript.

***James A. McLaughlin*** supervised the work of Anita Caron and Aleix Puig-Barbe.

***Katy Börner*** leads the HRA effort and contributed to writing.

***David Osumi-Sutherland*** built the initial prototypes of software described here, developed software specifications, supervised the work of Anita Caron and Aleix Puig-Barbe and is the primary author of this paper.

## Competing interests

The authors have no conflicts of interest to declare.

## Supplementary Figure

**Fig. S1.**
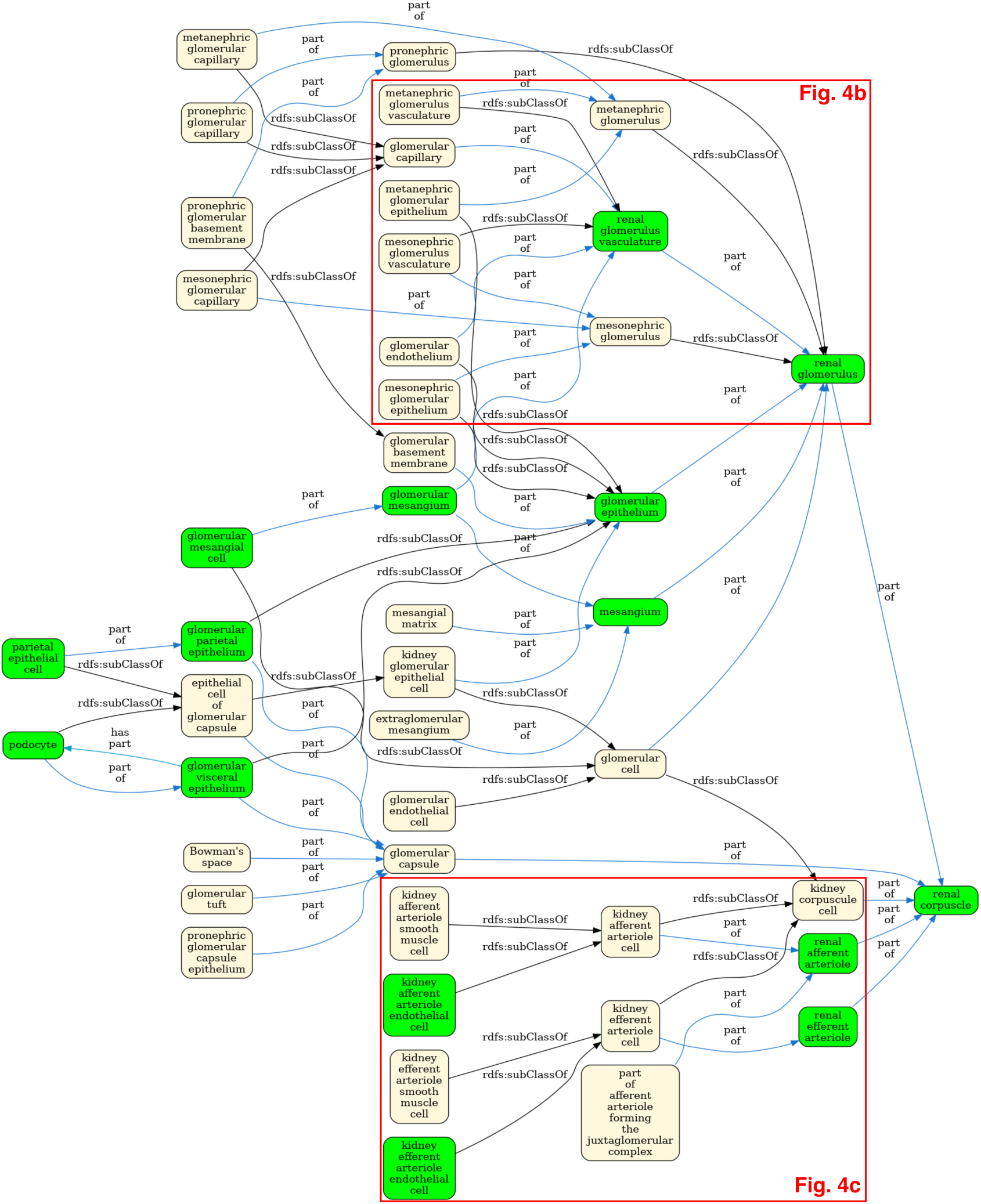
Uberon/CL ontology graph for the renal corpuscle with terms referenced in the HRA kidney ASCT+B table in green, including the renal corpuscle cell types illustrated in Fig. 4a. This illustrates the complexity of the Uberon graph compared to the needs of the HRA. Enlarged views of selected regions of the graph are provided in Fig. 4b and Fig. 4c.

## Notes

### Competing Interest Statement

The authors have declared no competing interest.

### Summary of Updates

We have revised the manuscript to improve clarity and strengthen the presentation. Specifically, we added further examples and explanations throughout the text to clarify key points, included an example SPARQL query in the Methods section, and revised the figures and expanded the figure legends to better support interpretation of the data and illustrate tool functionality.

## References

1. Wilkinson, M. D. et al. The FAIR Guiding Principles for scientific data management and stewardship. Sci. Data 3, 160018 (2016).

2. CZI Single-Cell Biology Program et al. CZ CELL×GENE Discover: A single-cell data platform for scalable exploration, analysis and modeling of aggregated data. bioRxiv 2023.10.30.563174 (2023) doi:10.1101/2023.10.30.563174.

3. Mi, H. et al. Protocol Update for large-scale genome and gene function analysis with the PANTHER classification system (v.14.0). Nat. Protoc. 14, 703–721 (2019).

4. Huntley, R. P. et al. A method for increasing expressivity of Gene Ontology annotations using a compositional approach. BMC Bioinformatics 15, 155 (2014).

5. Hitzler, P., Krötzsch, M., Parsia, B., Patel-Schneider - W3C …, P. F. & 2009. OWL 2 web ontology language primer. w3.org (2009).

6. Court, R. et al. Virtual Fly Brain-An interactive atlas of the Drosophila nervous system. Front. Physiol. 14, 1076533 (2023).

7. Rector, A. L. Modularisation of Domain Ontologies Implemented in Description Logics and Related Formalisms Including OWL. in Proceedings of the 2Nd International Conference on Knowledge Capture 121–128 (ACM, New York, NY, USA, 2003).

8. Osumi-Sutherland, D., Courtot, M., Balhoff, J. P. & Mungall, C. Dead simple OWL design patterns. J. Biomed. Semantics 8, 18 (2017).

9. Hill, D. P. et al. Dovetailing biology and chemistry: integrating the Gene Ontology with the ChEBI chemical ontology. BMC Genomics 14, 513 (2013).

10. Diehl, A. D. et al. The cell ontology 2016: Enhanced content, modularization, and ontology interoperability. J. Anim. Sci. Biotechnol. 7, (2016).

11. Costa, M., Reeve, S., Grumbling, G. & Osumi-Sutherland, D. The Drosophila anatomy ontology. J. Biomed. Semantics 4, 32 (2013).

12. Haendel, M. A. et al. Unification of multi-species vertebrate anatomy ontologies for comparative biology in Uberon. J. Biomed. Semantics 5, 21 (2014).

13. Gene Ontology Consortium. The Gene Ontology resource: enriching a GOld mine. Nucleic Acids Res. 49, D325–D334 (2021).

14. Gillespie, T. H., Tripathy, S., Sy, M. F., Martone, M. E. & Hill, S. L. The Neuron Phenotype Ontology: A FAIR approach to proposing and classifying neuronal types. bioRxiv 2020.09.01.278879 (2020) doi:10.1101/2020.09.01.278879.

15. Mungall, C. J., Koehler, S., Robinson, P., Holmes, I. & Haendel, M. k-BOOM: A Bayesian approach to ontology structure inference, with applications in disease ontology construction. bioRxiv (2016) doi:10.1101/048843.

16. Ding, S.-L. et al. Comprehensive cellular-resolution atlas of the adult human brain. J. Comp. Neurol. 524, 3127–3481 (2016).

17. Sikkema, L. et al. An integrated cell atlas of the lung in health and disease. Nat. Med. 29, 1563– 1577 (2023).

18. Bakken, T. E. et al. Comparative cellular analysis of motor cortex in human, marmoset and mouse. Nature 598, 111–119 (2021).

19. Yao, Z. et al. A high-resolution transcriptomic and spatial atlas of cell types in the whole mouse brain. Nature 624, 317–332 (2023).

20. Dorkenwald, S. et al. FlyWire: online community for whole-brain connectomics. Nat. Methods 19, 119–128 (2022).

21. Börner, K. et al. Anatomical structures, cell types and biomarkers of the Human Reference Atlas. Nat. Cell Biol. 23, 1117–1128 (2021).

22. Haniffa, M. et al. A roadmap for the Human Developmental Cell Atlas. Nature 597, 196–205 (2021).

23. Kong, Y. & Börner, K. Publication, funding, and experimental data in support of Human Reference Atlas construction and usage. Sci. Data 11, 574 (2024).

24. Börner, K. et al. Human BioMolecular Atlas Program (HuBMAP): 3D Human Reference Atlas Construction and Usage. Bioinformatics (2024).

25. Rosse, C. & Mejino, J. L. V., Jr. The foundational model of anatomy ontology. in Anatomy Ontologies for Bioinformatics 59–117 (Springer London, London, 2008).

26. Seal, R. L. et al. Genenames.Org: The HGNC resources in 2023. Nucleic Acids Res. 51, D1003– D1009 (2023).

27. Quardokus, E. M., Record, E. & Herr, B. W., II. SOP: Authoring Anatomical Structures, Cell Types and Biomarkers (ASCT+B) tables. Preprint at 10.5281/ZENODO.7382751 (2022).

28. Balhoff, J. & Curtis, C. k. INCATools/ubergraph: Release 2021-03-26. (Zenodo, 2021). doi:10.5281/ZENODO.4641309.

29. Balhoff, J. et al. Ubergraph: integrating OBO ontologies into a unified semantic graph. CEUR Workshop Proceedings (2022).

30. Börner, K. et al. Human BioMolecular Atlas Program (HuBMAP): 3D Human Reference Atlas construction and usage. Nat. Methods 22, 845–860 (2025).

31. Jain, S., et al. Anatomical Structures, Cell Types, plus biomarkers (ASCT+B) table for kidney, v1.5. HuBMAP 10.48539/HBM453.MFSJ.554 (2024).

32. Jackson, R. et al. OBO Foundry in 2021: operationalizing open data principles to evaluate ontologies. Database (Oxford) 2021, (2021).

33. Sridhar, M. S. Anatomy of cornea and ocular surface. Indian J. Ophthalmol. 66, 190–194 (2018).

34. Geven, E. J. W. & Klaren, P. H. M. The teleost head kidney: Integrating thyroid and immune signalling. Dev. Comp. Immunol. 66, 73–83 (2017).

35. Bajema, R. 2D renal corpuscle functional tissue unit (FTU) for Kidney, v1.2. (2023) doi:10.48539/HBM778.SHGJ.294.

36. Kruse, A., Schey, K. & Curcio, C. Anatomical Structures, Cell Types, plus biomarkers (ASCT+B) table for eye v1.0. HuBMAP 10.48539/HBM369.RBVN.935 (2021).

37. Nakamura, H., Katahira, T., Matsunaga, E. & Sato, T. Isthmus organizer for midbrain and hindbrain development. Brain Res. Brain Res. Rev. 49, 120–126 (2005).

38. Wexler, E. M. & Geschwind, D. H. Out FOXing Parkinson disease: where development meets neurodegeneration. PLoS Biol. 5, e334 (2007).

39. Rothstein, M., Bhattacharya, D. & Simoes-Costa, M. The molecular basis of neural crest axial identity. Dev. Biol. 444, S170–S180 (2018).

40. Martik, M. L. & Bronner, M. E. Riding the crest to get a head: neural crest evolution in vertebrates. Nat. Rev. Neurosci. 22, 616–626 (2021).

41. Jackson, R. C. et al. ROBOT: A tool for automating ontology workflows. BMC Bioinformatics 20, 407 (2019).

42. OpenStax AnatPhys fig.25.8 - The Kidney - English labels.

43. Pryhuber, G. Anatomical structures, cell types, plus biomarkers (ASCT+B) table for lung, v1.2. HuBMAP 10.48539/HBM496.MNTH.952 (2023).

44. Pryhuber, G. Anatomical structures, cell types, plus biomarkers (ASCT+B) table for lung, v1.5. HuBMAP 10.48539/HBM827.NKZG.839 (2024).

45. Miihkinen, M., Chu, Y., Vakkilainen, S., Akimov, Y. & Aittokallio, T. Estimating the completeness of large-scale single-cell sequencing projects. bioRxiv (2025) doi:10.1101/2025.01.08.631769.

46. Gray, J. I. & Farber, D. L. Tissue-resident immune cells in humans. Annu. Rev. Immunol. 40, 195– 220 (2022).

47. Osumi-Sutherland, D. et al. Cell type ontologies of the Human Cell Atlas. Nat. Cell Biol. 23, 1129– 1135 (2021).

48. Herr, B. W., 2nd et al. Specimen, biological structure, and spatial ontologies in support of a Human Reference Atlas. Sci. Data 10, 171 (2023).

